# A data-driven approach to quantifying meal characteristics influencing energy intake

**DOI:** 10.1101/2022.05.08.491083

**Authors:** Tera L. Fazzino, Amber B. Courville, Juen Guo, Kevin D. Hall

**Affiliations:** Department of Psychology, University of Kansas; Cofrin Logan Center for Addiction Research and Treatment, University of Kansas; National Institute of Diabetes and Digestive and Kidney Diseases, NIH, Bethesda, MD

## Abstract

We used a data-driven approach to determine the influence of energy density, hyper-palatability, protein content, and eating rate on *ad libitum* non-beverage energy intake during 2733 meals consumed by 35 inpatient adults who participated in two 28-day feeding studies. All four meal characteristics significantly contributed to energy intake, but their relative importance varied by the prevailing dietary pattern according to macronutrient composition and degree of processing.

## Main

Diets for the prevention and treatment of obesity are often informed by theories about food characteristics believed to support spontaneous reductions in energy intake without inducing hunger. Previous research relying primarily on self-reported food intake or single laboratory eating occasions suggests that energy density (ED), eating rate (ERate), protein content (%Prot), and proportion of hyper-palatable foods (%HPF) are potentially important meal characteristics influencing energy intake.^1-4^ We used a data-driven approach to determine the relative influence of these factors on *ad libitum* intake of 2733 meals consumed by 35 inpatient adults (**Table S1**) who participated in two feeding studies lasting 28 continuous days comparing minimally processed diets that varied in carbohydrate versus fat^5^ or diets with a balanced proportion of carbohydrate and fat that varied in ultra-processed versus minimally processed foods. ^6^

Using all the meal data available across studies, the **Table** shows that ED, ERate, %Prot, and %HPF significantly positively contributed to non-beverage energy intake within a linear mixed effects model accounting for whether the meal was breakfast, lunch, or dinner. ED and ERate had the strongest standardized effects, followed by %Prot and %HPF. Inclusion of all model covariates provided a substantially better fit to the data even after penalization for additional parameters according to Bayesian or Akaike’s Information Criteria. The **Figure** illustrates that modeled meal energy intake was highly correlated with actual intake (r=0.75; p<0.0001) with a mean absolute model error of ∼170 kcal which was ∼23% of the mean meal size. Neither previous meal intake of protein (β=0.27±0.57 kcal/g; p=0.63) nor energy (β=0.023±0.033 kcal/kcal; p=0.48) significantly affected subsequent meal energy intake using all meal data without accounting for dietary pattern.

**Table.** Linear Mixed Models of Meal Energy Intake.

| **Table. Linear Mixed Models of Meal Energy Intake** |  |  |  |  |
| --- | --- | --- | --- | --- |
|  | Standardized Beta | Std Error | p value | 95% CI |
| **All Diets (2733 meals; N=35)** |  |  |  |  |
| ED | 90.6 | 6.03 | <.0001 | 78.8 to 102.4 |
| ERate | 51.8 | 5.61 | <.0001 | 40.8 to 62.8 |
| %Prot | 37.8 | 4.73 | <.0001 | 28.5 to 47.1 |
| %HPF | 22.0 | 5.84 | .0002 | 10.6 to 33.5 |
| **Low Fat Diet (548 meals; N=20)** |  |  |  |  |
| ED | 46.8 | 12.35 | <.0001 | 22.5 to 71.1 |
| ERate | 61.8 | 10.72 | <.0001 | 40.7 to 82.8 |
| %Prot | -1.2 | 8.85 | 0.888 | -18.6 to 16.1 |
| %HPF | 8.8 | 14.67 | 0.550 | -20.1 to 37.6 |
| **Low Carbohydrate Diet (550 meals; N=20)** |  |  |  |  |
| ED | 75.9 | 10.73 | <.0001 | 54.8 to 96.9 |
| ERate | 127.5 | 12.36 | <.0001 | 103.2 to 151.7 |
| %Prot | 2.8 | 10.09 | 0.782 | -17 to 22.6 |
| %HPF | -4.1 | 9.22 | 0.659 | -22.2 to 14 |
| **Ultra-processed Diet (796 meals; N=20)** |  |  |  |  |
| ED | 15.0 | 13.55 | 0.269 | -11.6 to 41.6 |
| ERate | 32.0 | 10.39 | 0.002 | 11.6 to 52.4 |
| %Prot | 44.3 | 7.74 | <.0001 | 29.1 to 59.5 |
| %HPF | 54.8 | 12.70 | <.0001 | 29.9 to 79.7 |
| **Unprocessed Diet (839 meals; N=20)** |  |  |  |  |
| ED | 93.7 | 8.00 | <.0001 | 78 to 109.4 |
| ERate | 51.8 | 9.26 | <.0001 | 33.7 to 70 |
| %Prot | 48.3 | 8.98 | <.0001 | 30.7 to 65.9 |
| %HPF | 6.9 | 9.05 | 0.447 | -10.9 to 24.6 |
| *Note.* CI= confidence interval; %HPF= percent energys from hyper-palatable foods; ED= energy density in kcal per gram; %Prot= percent energys from protein; ERate= eating rate in grams per minute. Meal as a factor (breakfast/lunch/dinner) was also included as a covariate in all models. |  |  |  |  |

**Figure.**
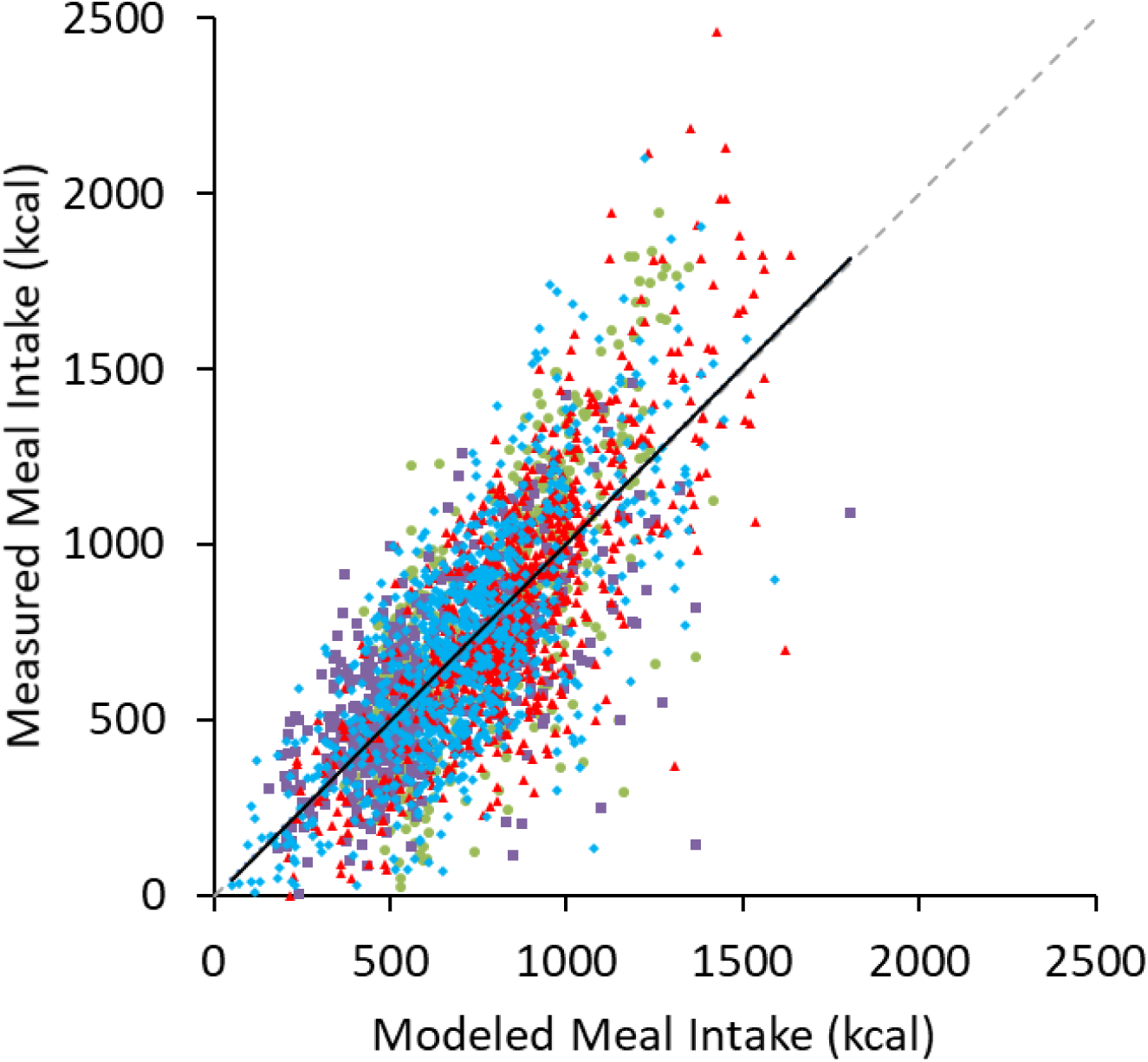
Modeled versus measured meal energy intake. A linear mixed-model relating meal energy density, eating rate, proportion of hyperpalatable foods, and protein content was used to predict *ad libitum* energy intake of 2733 meals consumed by 35 inpatient adults exposed to low carbohydrate diets (green circles), low fat diets (purple squares), diets high in ultra-processed food (red triangles) and diets high in minimally processed foods (blue diamonds). The line of identity is indicated as the dashed gray diagonal line and the solid black line is the best fit regression line.

Separating the data by dietary pattern revealed that ERate was a significant positive contributor to meal energy intake in all diets (Table). ED was a significant positive contributor to energy intake in all diets except the ultra-processed diet where %HPF positively contributed to energy intake, but ED was highly correlated with %HPF (r=0.768; **Table S2**) making it potentially difficult to disentangle the relative contribution of ED and %HPF within this dietary pattern. During the unprocessed diet, all factors except %HPF were significant positive contributors to meal energy intake. During both the low fat and low carbohydrate diets, neither %Prot nor %HPF were significant contributors to energy intake.

Previous intake significantly affected meal energy intake during both the low fat and low carbohydrate diets, with previous meal protein intake positively influencing subsequent meal energy intake (low fat β=6.5±1.1 kcal/g; p<0.0001, low carbohydrate β=5.6±1.9 kcal/g; p=0.003) whereas previous meal energy intake negatively influenced subsequent meal energy intake (low fat β=-0.30±0.06 kcal/g; p<0.0001 and low carbohydrate β=-0.27±0.08 kcal/g; p=0.001). During the unprocessed diet, previous meal energy intake negatively influenced subsequent meal energy intake (β=-0.22±0.06 kcal/kcal; p=0.0008) whereas previous meal protein intake had no significant effect (β=-0.2±1.0 kcal/g; p=0.85). During the ultra-processed diet, previous meal protein intake negatively influenced subsequent meal energy intake (β=-3.1±1.3 kcal/g; p=0.02) whereas previous meal energy intake had no significant effect (β=0.035±0.088 kcal/kcal; p=0.69). Previous meal intake of energy and protein were highly correlated across all diets (r>0.82).

The primary outcomes of the original studies found that the unprocessed diet resulted in ∼500 kcal/d less mean daily intake compared to the ultra-processed diet^6^ and the low fat diet resulted in ∼700 kcal/d less mean daily intake than the low carbohydrate diet.^5^ **Table S3** presents mediation analyses showing that the effect of unprocessed versus ultra-processed diets on non-beverage meal energy intake was significantly positively mediated by ED and

%HPF (ED 45.1±13.6%; p=0.001 and %HPF 41.9±6.3%; p<0.0001). **Table S4** shows that these factors also positively mediating the effect of low fat versus low carbohydrate diets (ED 24.4±5.5%; p<0.0001 and %HPF 14.0±4.0%; p=0.004). ERate significantly negatively mediated the association between diet condition and meal energy intake in both studies because mean non-beverage meal eating rate was lower in both the ultra-processed and low carbohydrate diets (p values <0.0001). %Prot significantly negatively mediated the effect of ultra-processed versus unprocessed diets because the mean protein content of the ultra-processed meals was slightly lower than the unprocessed meals but %Prot did not significantly mediate the effect of low fat versus low carbohydrate diets where protein content was more closely matched.

In summary, the meal characteristics ED, ERate, %HPF, and %Prot were all important for predicting energy intake, but their relative importance varied depending on dietary pattern. Overall, meals with greater energy density, comprised of more hyperpalatable foods with higher protein content that were eaten more rapidly resulted in greater energy intake within an eating occasion. Mediation analyses consistently identified energy density and hyperpalatable foods as positively mediating the effects of diet condition on meal energy intake. Future studies should prospectively test the impact of manipulating food and meal-based characteristics over time and determine whether energy intake spontaneously decreases by choosing meals with reduced energy density, lower protein content, and comprising fewer hyper-palatable foods that are eaten more slowly. Our findings may identify points of leverage for influencing energy intake in a manner that does not require effortful attention to energy intake on the part of the individual, which is a consistent challenge in obesity prevention and dietary adherence.

## Methods

This was a secondary analysis of data collected from two previous studies^5,6^ conducted at the Metabolic Clinical Research Unit at the NIH Clinical Center that were approved by the Institutional Review Board of the National Institute of Diabetes & Digestive & Kidney Diseases. All participants provided informed consent. Eligibility criteria were: 1) ages 18-50y; 2) BMI>18.5 kg/m^2^; and 3) weight stable (<5% change in past 6 months). See Supplementary Table S1 for baseline information on the participants. Both studies used a within-subjects, random-order, crossover design to expose participants to two diet conditions with 7-day rotating menus for 14 days each. Participants were instructed to not try to change their weight and eat as much or as little as they wanted. All foods were weighed to the nearest 0.1 grams before and after consumption and energy intake was calculated using ProNutra software.

Presented meals were analyzed at the individual food level without beverages. Hyper-palatable foods (HPF) were defined as being high in fat and sodium, high in fat and sugar, or high in carbohydrate and sodium using a standardized definition described previously.^4^ Percentage of presented meal energy from HPF (%HPF) and protein (%Prot) were used as predictor variables along with the meal energy density (ED) in kilocalories per gram and the meal eating rate (ERate) in grams of food consumed per minute. Linear mixed effects models with meal energy intake as the dependent variable included a random intercept for participant and specified an exchangeable correlation structure with meal type (breakfast, lunch, or dinner) as a covariate, along with %HPF, %Prot, ED, and ERate. In addition to within-meal analyses, we investigated whether protein and energy intake of the previous meal within each day influenced subsequent meal energy intake, given the hypothesis regarding the role of protein to increase satiety^2^. Finally, mediation analyses were conducted with each variable included as a mediator between diet condition and meal energy intake.

## Supplemental Information

### Diets and Participants

Adult participants were admitted as inpatients for a continuous 28-day trial, with each diet presented for 14 consecutive days in random order. Three meals per day were provided on a 7-day rotating menu and meals were categorized according to the NOVA categorization system to identify ultra-processed or minimally processed foods (ultra-processed vs. unprocessed diet study) or macronutrient content for carbohydrates and fat (low fat vs. low carbohydrate diet study). Twenty adults completed each study and 5 completed both studies for a total of 35 unique participants.

**Table S1.**
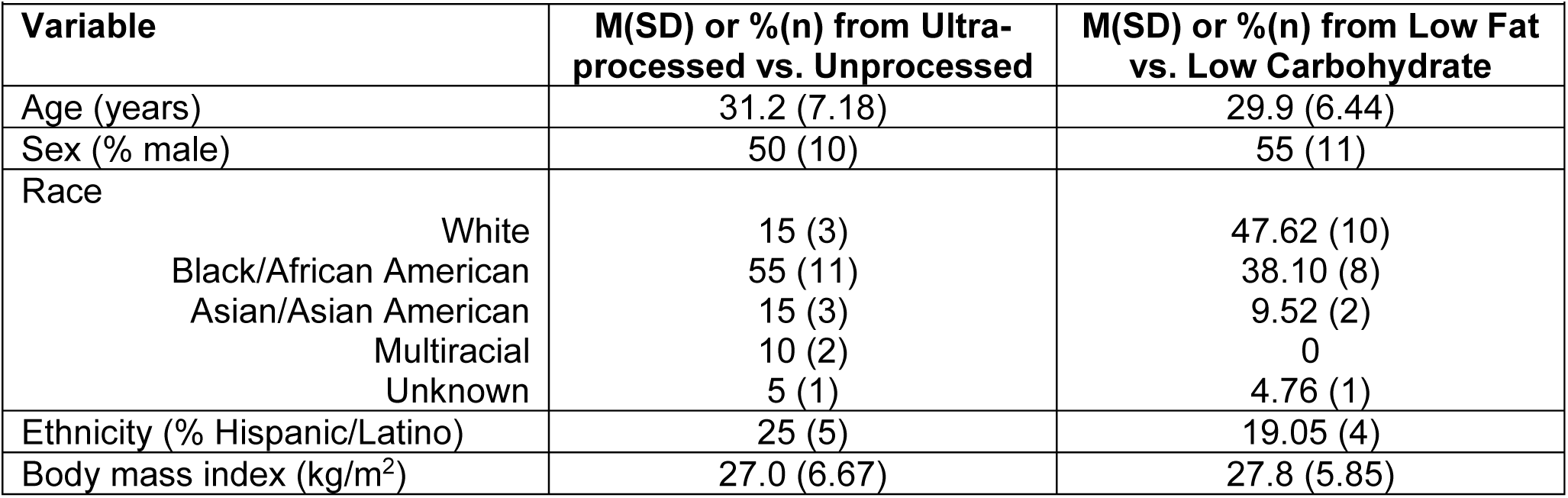
Participant Characteristics (N=35)

**Table S2.** Correlation Matrices.

| **Table S2. Correlation Matrices** |  |  |  |  |
| --- | --- | --- | --- | --- |
| **Ultra-processed Diet** |  |  |  |  |
| Variable | %HPF | ED | ERate | %Prot |
| %HPF | 1 |  |  |  |
| ED | 0.768 | 1 |  |  |
| ERate | -0.168 | -0.294 | 1 |  |
| %Prot | -0.150 | -0.257 | 0.085 | 1 |
| **Unprocessed Diet** |  |  |  |  |
| Variable | %HPF | ED | ERate | %Prot |
| %HPF | 1 |  |  |  |
| ED | 0.181 | 1 |  |  |
| ERate | 0.110 | -0.035 | 1 |  |
| %Prot | 0.111 | -0.153 | -0.009 | 1 |
| **Low Fat Diet** |  |  |  |  |
| Variable | %HPF | ED | ERate | %Prot |
| %HPF | 1 |  |  |  |
| ED | 0.411 | 1 |  |  |
| ERate | -0.162 | -0.303 | 1 |  |
| %Prot | -0.103 | -0.194 | 0.172 | 1 |
| **Low Carbohydrate Diet** |  |  |  |  |
| Variable | %HPF | ED | ERate | %Prot |
| %HPF | 1 |  |  |  |
| ED | 0.228 | 1 |  |  |
| ERate | 0.021 | -0.234 | 1 |  |
| %Prot | 0.065 | -0.067 | -0.088 | 1 |
| **All Diets Combined** |  |  |  |  |
| Variable | %HPF | ED | ERate | %Prot |
| %HPF | 1 |  |  |  |
| ED | 0.567 | 1 |  |  |
| ERate | -0.136 | -0.293 | 1 |  |
| %Prot | -0.022 | -0.093 | 0.035 | 1 |
| *Note.* %HPF= percent of presented meal energy from hyper-palatable foods; ED= energy density in kcal/g; %Prot= percent of presented meal energy from protein; ERate= meal eating rate in g/min |  |  |  |  |

**Table S3.**
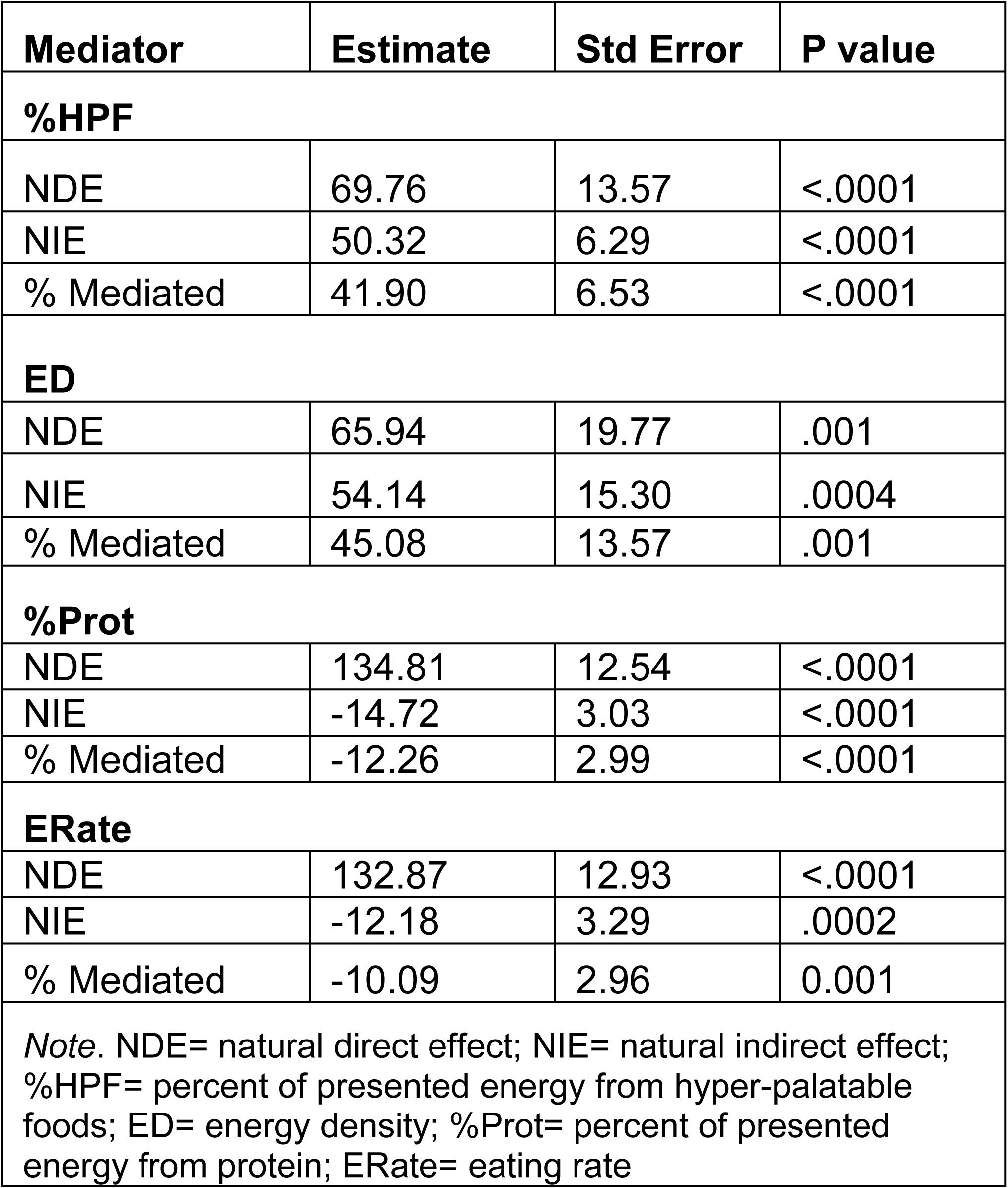
Mediation Analyses for the Ultra-processed vs. Unprocessed Diet Study.

**Table S4.** Mediation Analyses for the Low Fat vs. Low Carbohydrate Diet Study.

|  |  |  |  |
| --- | --- | --- | --- |
| **Mediator** | **Estimate** | **Std Error** | **P value** |
| **%HPF** |  |  |  |
| NDE | 159.34 | 12.92 | <.0001 |
| NIE | 25.91 | 7.22 | <.0001 |
| % Mediated | 13.99 | 3.97 | .0004 |
| **ED** |  |  |  |
| NDE | 140.14 | 14.48 | <.0001 |
| NIE | 45.12 | 9.79 | <.0001 |
| % Mediated | 24.35 | 5.45 | <.0001 |
| **%Prot** |  |  |  |
| NDE | 185.27 | 11.01 | <.0001 |
| NIE | -0.01 | 2.10 | .997 |
| % Mediated | -0.01 | 1.14 | .997 |
| **ERate** |  |  |  |
| NDE | 183.43 | 13.73 | <.0001 |
| NIE | -19.46 | 4.33 | <.0001 |
| % Mediated | -11.87 | 2.87 | <.0001 |
| *Note.* NDE= natural direct effect; NIE= natural indirect effect; %HPF= percent of presented energy from hyper-palatable foods; ED= energy density; %Prot= percent of presented energy from protein; ERate= eating rate |  |  |  |

## Notes

**Funding:** This work was supported by the Intramural Research Program of the National Institutes of Health, National Institute of Diabetes & Digestive & Kidney Diseases.

**Conflict of interest disclosure statement:** None of the authors have conflicts of interest

### Competing Interest Statement

The authors have declared no competing interest.

